# Student experiences with an interactive 3D immersive biotechnology simulation and its impact on motivational beliefs

**DOI:** 10.1101/2024.01.23.576860

**Authors:** Dan Spencer, Caitlin McKeown, David Tredwell, Benjamin Huckaby, Andrew Wiedner, Jacob T. Dums, Emily L. Cartwright, Colin M. Potts, Nathan Sudduth, Evan Brown, Phillips Albright, Arnav Jhala, Melissa C. Srougi

**Affiliations:** Digital Education and Learning Technology Applications, North Carolina State University, Raleigh, North Carolina, United States of America; Biotechnology Program, North Carolina State University, Raleigh, North Carolina, United States of America; Department of Computer Science, North Carolina State University, Raleigh, North Carolina, United States of America; Department of Molecular Biomedical Sciences, North Carolina State University, Raleigh, North Carolina, United States of America

## Abstract

The development and use of virtual laboratories to augment traditional in-person skills training continues to grow. Virtual labs have been implemented in a number of diverse educational settings, which have many purported benefits including their adaptability, accessibility, and repeatability. However, few studies have evaluated the impact of virtual laboratories outside of academic achievement and skills competencies, especially in biotechnology. In this study, an interdisciplinary team of content experts, video game researchers, instructional designers, and assessment experts developed a 3D immersive simulation designed to teach novice scientists the technical skills necessary to perform sterile mammalian cell culture technique. Unique to the simulation development process is the recreation of an immersive experience through the capture of details in the real-world lab where participants have the freedom of choice in their actions, while receiving immediate feedback on their technical skills as well as procedural execution. However, unlike an in-person laboratory course, students are able to iterate and practice their skills outside of class time and learn from their mistakes. Using a mixed-methods study design, over the course of two semesters we evaluated student attitudes of the simulation and their science motivational beliefs including self-efficacy and science identity after engaging with the simulation prior to the physical laboratory. Our results show that students’ science identity remained unchanged while their science self-efficacy increased. Furthermore, students had positive perceptions of the benefits of the virtual simulation. These data suggest that the virtual cell culture simulation can be a useful pedagogical training tool to bolster students’ motivational beliefs that is both accessible and easy to implement.

## Introduction

Laboratory training is a critical component to many STEM courses as it is the practical application of concepts and theory through hands-on, in-person experiences. The benefits of laboratory training are numerous as it affords students opportunities to practice and promote key analytical, problem-solving, and critical thinking skills. Moreover, it allows students to learn cutting-edge technical skills first-hand as practicing scientists while keeping pace with the ever evolving scientific landscape.

Recent advances in technology have introduced virtual elements to the classical physical laboratory training, which range from pre-recorded demonstration videos, interactive questions, and point and click simulations, to fully immersive virtual reality laboratories complete with virtual teaching assistants [1]. Both physical and virtual laboratory training allow students to learn the process of science, develop specific technical skill sets, and promote their conceptual understanding of the material. Virtual laboratory training, in particular, has provided new teaching tools and opportunities for student learning. Virtual labs can be adaptable and focus on elements that create the most challenge for students, such as those not readily observable [2]. When used alone, virtual gamification of labs increased student learning gains compared to traditional physical teaching methods specifically within biotechnology education [2]. Moreover, students reported higher confidence using and operating lab equipment [3], and the combination of both the physical and virtual learning modalities resulted in greater learning gains for students. However, not much is known about user motivational beliefs when exposed to virtual training. As students navigate the immersive realm of virtual laboratory training, their perceptions alter, giving way to deeper understanding of the potential benefits and applications of learning in the virtual environment. Therefore, identifying the motivational beliefs that surround simulation use as a training tool is critical in determining how students invest in the virtual experience. This is becoming increasingly important as virtual laboratory training increases in both academic and industry sectors.

In general, virtual laboratories have been positively received by students [4–7] and have garnered high satisfaction ratings from both students and instructors [3, 8]. Further, students have expressed beliefs that virtual laboratories promote learning [6, 7], increase their awareness [9] and attitudes towards a subject area [10], as well as simplify complex scientific processes [8]. However, virtual laboratories are often seen as a preparatory tool or complementary to in-person experiences by students and instructors [6, 7, 11]. The COVID-19 pandemic shifted this paradigm demonstrating that virtual learning can be implemented and effective as stand-alone skills training tools [1, 12].

Understanding students and instructors’ general perceptions are important for the development and integration of virtual simulations in the college classroom due to their impact on exploration/engagement [13]. However, it is also important to consider how successful laboratories are in reaching their learning goals. This includes not only course specific knowledge gains and achievement outcomes but also how these experiences aid students’ growth as scientists in terms of both their identity (aka science identity) and self-efficacy.

Carlone and Johnson (2007) describe science identity as comprising of three dimensions: 1) student knowledge of science content, skills, and practices (competence), 2) student ability to conduct science practices and demonstrate competence to others (performance), and 3) self-acknowledgement or acknowledgement by others that a student is a science person (recognition) [14, 15]. Science identity should be viewed as malleable and impacted by student experiences and interactions [16]. Importantly, increases in students’ science identity have been linked to a stronger commitment to a science career [17], persistence and retention in the sciences [15, 18], as well as increases in student motivation and community engagement [19]. Conversely, identity mismatch, whereby students experience uncertainty about how they fit in an academic environment, can result in withdrawal from academic pursuits and opportunities [14].

While numerous in-depth studies have shown that hands-on in person laboratory training fosters a strong science identity in students [20], very little work has investigated the impact of virtual laboratory training [3, 7]. In fact, research assessing the validity of virtual laboratory training in microbiology and biology settings has often focused on preferences, understanding, and value of the simulation. However, based on research on the factors that impact science identity development [15, 21] and common instructional approaches in designing of laboratory simulations [10, 22], it is reasonable to assume that student engagement with virtual laboratory environments has the potential to impact their science identity analogous to physical laboratory training. In particular, elements of virtual simulations such as 1) the use of active learning [23], 2) reduction or removal of barriers to access [10], and 3) reduction of risk and providing space for students to fail [24] all have the potential to increase students’ science identity by bolstering competence, increasing performance, and promoting self-acknowledgement [25].

An important motivational belief that cultivates an individual’s science identity is self-efficacy [16, 21, 26, 27]. In the current study context, self-efficacy is viewed as a student’s self-reported confidence in understanding and using [biology] in their lives [28]. It is viewed as multidimensional in that it not only encapsulates confidence in methods, but also generalization and application of concepts and skills [28]. Self-efficacy, rooted in social cognitive theory, views achievement as based on various interactions between behavior, environment, and personal factors [26, 29]. Of note, self-efficacy in STEM fields is correlated with academic achievement [30], task persistence [31, 32], motivation [33, 34], and resilience [28, 35]. Recent studies examining large cohorts of undergraduate STEM graduates reported that both self-efficacy and science identity were critical and universal attributes of students committing to STEM careers [14]. Therefore, these studies underscore the significance of motivational beliefs in the evaluation of novel virtual STEM pedagogical training tools as they are becoming more commonplace in the digital era. Nevertheless, similar to investigations of the development of student science identity, very little work has been conducted on the impact of virtual laboratories on student self-efficacy [36]. Based on the conceptual understanding of the growth and formation of efficacy related beliefs, virtual laboratory training tools could therefore potentially impact student self-efficacy. This is due to the fact that the virtual environment affords risk taking by reducing the fear of failure, promoting goal achievement, increasing positive emotion, and having a direct translation to real-world applications [3].

Currently, there are few studies that evaluate the impact of biotechnology virtual laboratories outside of academic achievement and skills competencies [3]. Therefore, in the present study we not only investigate student perceptions of an immersive and interactive virtual laboratory simulation in a dual-enrollment undergraduate/graduate biotechnology classroom, we also examine student attitudes towards the simulation and potential changes in students’ science beliefs (self-efficacy and science identity) following its usage as a training tool. Specifically, this study addresses the following research questions:

1. What were students’ perceptions of the interactive virtual cell culture simulation?
2. How were students’ motivational beliefs (self-efficacy and science identity) impacted as a result of engaging with the interactive virtual cell culture simulation over time?

## Methods

### Virtual cell culture simulation description

The overarching goal of the project was to create an immersive and interactive, online lab experience by providing a configurable and virtual lab environment where students could evaluate their decisions based on immediate feedback. While many off-the-shelf virtual lab programs were available [37], they lacked the right combination of realism, customizability, and decision-making opportunities desired by the course biotechnology researchers and instructional design development team. Finally, the researchers wanted a virtual laboratory that was both based on the laboratory environment at their institution and also fully customizable for the widest range of experiments and experience levels.

The virtual simulation is a single-page web application that has a baseline laboratory model with commonly used items, as well as an array of optionally loadable items. The experience is available on a broad range of devices (i.e. tablets, laptops, cellular phones), with additional supplemental materials and activities that utilize alternative technologies, such as video.

The laboratory space itself was created based on real laboratory spaces, while also representing an ideal case in terms of space and budget conditions. Various floor plans, layouts, and designs were created and considered before ultimately deciding on a laboratory space that was realistic, but still generic.

Within the virtual laboratory environment students can navigate around the space, but are mostly limited to actions at different stations. A protocol is provided and each station hosts specific equipment and objects, but students are allowed to navigate freely between them and perform activities non-linearly.

To achieve this non-linear navigation within the simulation, students’ progress through the experiment by interacting with the equipment and objects at each station. By selecting objects alone or in various combinations, they can perform actions such as mixing and measuring solutions, changing settings on a device, or potentially introducing contamination. This information is stored per object, which serves as a system for saving and loading as well as measuring success and failure for procedure steps.

### Study Context/Course Description

The study took place within the course Manipulation of Recombinant DNA at an R1 research university. This is a full semester 4 credit hour upper-level course consisting of 2 h lecture and 5 h laboratory periods each week. The course enrolls both undergraduate and graduate students focusing on the theory and practice behind recombinant DNA cloning and screening, protein expression, and experimental design using a mammalian cell culture model system. The prerequisites for the course were two semesters of general chemistry and two semesters of organic chemistry. Three sections of the course were taught in both the Fall and Spring semesters by life sciences PhD holding faculty and teaching postdoctoral fellows. The lab simulation was used to supplement laboratory instruction of the course content and hands-on laboratories related to experimental design, mammalian sterile cell culture technique, cell plating, and transfection.

### Laboratory Simulation on Mammalian Cell Culture Sterile Technique and Implementation

The lab simulation was designed using the same learning outcomes as was embedded in the course. Course expectations and assessment methods were detailed in the syllabus, as were the learning outcomes. *Upon completion of the laboratory simulation, students should be able to:*

1. Practice sterile cell culture technique using a biosafety cabinet.
2. Passage mammalian cells using sterile technique.
3. Count cells using an automated hemocytometer and apply dilution calculations.

The lab simulation was used prior to students performing the exact same laboratory procedure in the physical laboratory. During their scheduled laboratory period, students used their electronic laboratory notebooks to access the link to the virtual simulation and went through the steps of the procedure virtually in the simulation. Teaching assistants discussed the simulation platform briefly including an overview of the interface, how to change location, and how to manipulate objects prior to beginning. Students were also given a Frequently Asked Questions document and a short video tutorial on the lab simulation ahead of time to further familiarize themselves with the simulation platform. Teaching assistants were on hand to help students should they require assistance. Students took an average of 1.5 hours to complete the lab simulation. Upon completion, they were directed to the cell culture facility to perform components of the identical laboratory procedure in-person. After completion of both the virtual and physical laboratory on sterile cell culture technique, students were required to complete a laboratory notebook entry and a short reflection to assess their learning.

### Instructional strategy and approach

Development was guided by priorities that were established early on in the analysis and design stages of the project. Specifically, the researchers wanted to emphasize critical thinking and diagnostic skills, so it was important to allow students to make mistakes, or even fail, in the course of completing the experiment. One key goal was to provide a virtual environment where students could practice their laboratory skills in a low-risk, time effective way, without concerns about contamination, incubation time, laboratory availability, or consumption of resources.

The researchers wanted a virtual laboratory that was as customizable and open-ended as a physical laboratory, so emphasis was placed on allowing opportunities for decision-making and promoting configurability of the lab environment. Process fidelity was also an important priority for the simulation, so it was designed to provide a first-person perspective and use photo-realistic imagery and environments. The simulation employs an instructional strategy that combines intrinsically programmed discovery with meaningful reception learning [22, 38]. Students demonstrate their understanding of the concepts by applying and executing them in the laboratory environment. While the virtual laboratory is programmed to allow for free movement and completion of steps, it also includes pre-programmed feedback and outcomes based on expected behaviors, common mistakes, and pre-established parameters. Students are further guided through the experience through the use of visual cues and prompts. For example, students receive real-time feedback on their technique through a risk-meter that is prominently displayed in the user interface. The color and bar length on the risk meter will change if a student engages in behavior that could introduce contamination into their sample.

### Participants

Data used for the current study were taken from an internal course redesign project during fall 2022 and spring 2023 semesters of the same academic year. A total of 179 (104 Fall, 75 Spring) adult students participated in the assessment of the redesign, with 152 (87 Fall, 65 Spring) participants agreeing to share their data for research purposes. Participants completed online surveys (Qualtrics Survey Software) at three different time points: 1) the week prior to engaging with the simulation during lab (n = 87 Fall, n = 65 Spring), 2) the week following the lab (n = 87 Fall, n = 64 Spring), and 3) 4 weeks following the lab which was at the end of semester (n = 73 Fall, n = 61 Spring). The NC State University Institutional Review Board (IRB) approved this protocol (#24300) under an exempt review.

### Qualitative Methods

#### General Perceptions of the Virtual Interactive Cell Culture Simulation

During the post-lab and end of semester survey, participants rated their general perceptions of the simulation using five items. Items required participants to rate on a five-point Likert scale from 1 (strongly disagree) to 5 (strongly agree) the extent to which the simulation 1) increased their engagement, 2) required them to think critically, 3) helped them make connections between prior and new knowledge, 4) increased their understanding of the importance of sterile mammalian cell culture technique, and 5) provided real-world applications. Responses were collated across items to form a single score that was used to categorize participants into different groups based on their overall perceptions of the simulation during analysis.

#### Open-ended feedback

Two open-ended questions were provided in the end of semester survey to allow participants to provide feedback on 1) the most helpful elements of the simulation and 2) ways the simulation could be improved to help students learn better.

#### Biotechnology self-efficacy

A measure of biotechnology self-efficacy was created by the researchers based on the work of Baldwin et al. (1999) [28]. During each survey participants were asked to rate their beliefs about their ability to complete eight specific biotechnology tasks on a 4-point Likert scale from 1 (Not at all confident) to 4 (totally confident). Reliability testing using Cronbach’s alpha showed the scale to be reliable across time points (**S1 Table**).

#### Science identity

Adapted from Chemers et al. (2011), participants were asked during pre-lab and end of semester surveys to think about how much they thought being a scientist is part of who they are [17]. Participants responded to six items on a 5-point Likert scale from 1 (strongly disagree) to 5 (strongly agree). Reliability testing using Cronbach’s alpha showed the scale to be reliable at both pre-lab and end of semester surveys (**S1 Table**).

### Analyses

Descriptive statistics and paired samples t-tests were used to explore participants’ perceptions of the interactive virtual cell culture simulation. Further, thematic analysis was undertaken to understand participants’ open-ended feedback relating to the most helpful features and areas to improve the simulation. Two members of the research team engaged in independent open coding resulting in themes emerging directly from the data. A subset of responses was chosen for the initial coding. Multiple codes could correspond to a single response. Following this, team members met to resolve discrepancies and discuss general themes. This process was repeated until themes were confirmed. As a final step, coders reviewed the data to confirm its alignment with the final list of themes. To increase the validity of the qualitative process, two validation strategies were utilized: memoing and peer debriefing [39]. Coders engaged in memoing by noting ideas and documenting processes/decisions taken during the coding process. To reduce bias, peer debriefing was conducted with the remaining members of the research team through targeted conversations on the coders approach and interpretations of the qualitative data. During this process, the research team set the threshold that at least 10 percent of student responses must include a topic for it to be considered a theme.

To explore how participants’ motivational beliefs (self-efficacy and science identity) were impacted as a result of engaging with the interactive virtual cell culture simulation, participants were broken into three groups using a tertile split of data relating to their general perceptions of the simulation. Cut points used to create the groups were based on the five point Likert scale used for the measure. Participants were placed in the “low group” if they averaged below a three out of five on the scale (n = 44), in the “moderate” group if they averaged between a three and a four (n = 48), and in the “high” group if they averaged a four or above. Quantitative analyses were then conducted to understand changes in student motivation over time by simulation perception. A mixed 3x3 (time x group) ANOVA was used for biotechnology self-efficacy and a 2x3 ANOVA for science identity. Data met all test assumptions [40].

## Results

### Course Context and Simulation Implementation

Results are presented from surveys from 152 consenting students enrolled in an interdisciplinary dual enrollment upper-level molecular biotechnology lecture and laboratory course. Three sections of the course are offered every Fall and Spring semesters and include 2 h of lecture and 5 h of laboratory once per week. Students participating in this course were from numerous majors including but not limited to chemical engineering, physiology, biological sciences, microbiology and others (**Table 1**). Participant demographics are detailed in (**Table 2**). Based on information obtained from university student records, the majority of participants were predominantly female 62.25% (n=94) and 37.75% (n=57) male. Of the total, 74.17% (n=112) were traditional in age (i.e. 18-24 years) and 25.83% (n=39) were non-traditional in age (i.e. >24 years). Sixty percent (60.27%; n=91) of participants were white, 12.58% (n=19) were Asian, 6.62% (n=10) were Hispanic or LatinX, 5.96% (n=9) were Black or African American, 4.64% (n=7) were two or more races, 0.66% (n=1) were Pacific Islander, and 9.27% were classified as Non-resident Alien (**Table 2**). Sixty-one percent (61.59%; n=93) of participants were undergraduates, 36.42% (n=55) were graduate level, and 1.99% (n=3) were other.

**Table 1.** Course Enrollment Information by Major and Level.

| COLLEGE | MAJOR | NUMBER OF GRAD/UNDERGRAD | TOTAL STUDENTS |
| --- | --- | --- | --- |
| AGRICULTURE AND LIFE SCIENCES | Animal Science | 1/1 | 2 |
|  | Biochemistry | 1/6 | 7 |
|  | Biological Engineering | 0/1 | 1 |
|  | Crop and Soil Sciences | 0/3 | 3 |
|  | Entomology | 1/0 | 1 |
|  | Horticultural Science | 0/2 | 2 |
|  | Plant Biology | 0/2 | 2 |
|  | Plant Pathology | 2/0 | 2 |
|  | Soil Science | 1/0 | 1 |
| ENGINEERING | Bioinformatics | 1/0 | 1 |
|  | Biomedical Engineering | 0/5 | 5 |
|  | Chemical Engineering | 7/25 | 32 |
|  | Civil Engineering | 1/0 | 1 |
|  | Industrial Engineering | 1/0 | 1 |
|  | Mechanical Engineering | 0/2 | 2 |
| SCIENCES | Biological Sciences | 1/23 | 23 |
|  | Biology | 1/0 | 1 |
|  | Chemistry | 5/0 | 5 |
|  | Genetics | 0/7 | 7 |
|  | Microbial Biotechnology | 7/0 | 7 |
|  | Microbiology | 2/13 | 15 |
|  | Physics | 1/0 | 1 |
|  | Physiology | 10/na | 10 |
|  | Zoology | 0/2 | 2 |
| **VETERINARY MEDICINE** | Comparative Biomedical Sciences | 7/0 | 7 |
| **NATURAL RESOURCES** | Environmental Sciences | 0/1 | 1 |
|  | Forestry and Environmental Sciences | 2/0 | 2 |
| **TEXTILES** | Fiber and Polymer Sciences | 4/0 | 4 |
| **HUMANITIES AND SOCIAL SCIENCES** | Psychology | 0/1 | 1 |
| **NONE** | Post baccalaureate studies | 0/2 | 2 |

**Table 2.** Student Demographic Information.

| **Subcategory** | **%** |
| --- | --- |
| Female | 62.25 |
| Male | 37.75 |

**Race**
| **Subcategory** | **%** |
| --- | --- |
| Asian | 12.58 |
| Black or African American | 5.96 |
| Hispanic of any race | 6.62 |
| Non-resident alien | 9.27 |
| Pacific Islander | 0.66 |
| Two or more races | 4.64 |
| White | 60.27 |

<sup>a</sup>One student was missing from the data.

A majority of students sampled for the current study had taken at least one biotechnology course previously (30.20% no experience, 37.58% 1 course, 28.86% 2-4 courses, 3.36% 5+ courses). A majority (84.87%) also indicated that they were somewhat or extremely comfortable with molecular biology (0.66% extremely uncomfortable, 5.26% somewhat uncomfortable, 9.21% neither comfortable nor uncomfortable). However, less than half of participants (44.08%) reported feeling (somewhat or extremely) comfortable with the technology used to deliver the simulation prior to using it (41.45% neither comfortable nor uncomfortable; 12.50% somewhat uncomfortable, 1.97% extremely uncomfortable).

A 3D immersive virtual laboratory simulation that focused on the technical skills required to perform sterile mammalian cell culture technique was developed. The simulation afforded students the ability to make independent decisions in the virtual environment that directly impacted experimental outcomes while receiving real-time feedback on their technique through an integrated risk-meter (**Fig 1A**). The simulation was modeled on real laboratory spaces and students followed protocol steps in an electronic laboratory notebook that was integrated into the simulation (**Figs 1A and B**). The simulation was implemented during the unit of the course, which discusses sterile mammalian cell culture technique and transfection. During this unit, students in all sections were assigned to complete the virtual interactive simulation during laboratory time. Teaching assistants (TAs) went over the simulation interface with students and provided a FAQ guide prior to engaging in the simulation. Students completed the simulation on average in ∼1.5 h. Once students completed the simulation, they performed the same experiment in the physical lab space. TAs were available during the lab to answer questions related to the simulation technology and experimental technique.

**Fig 1.**
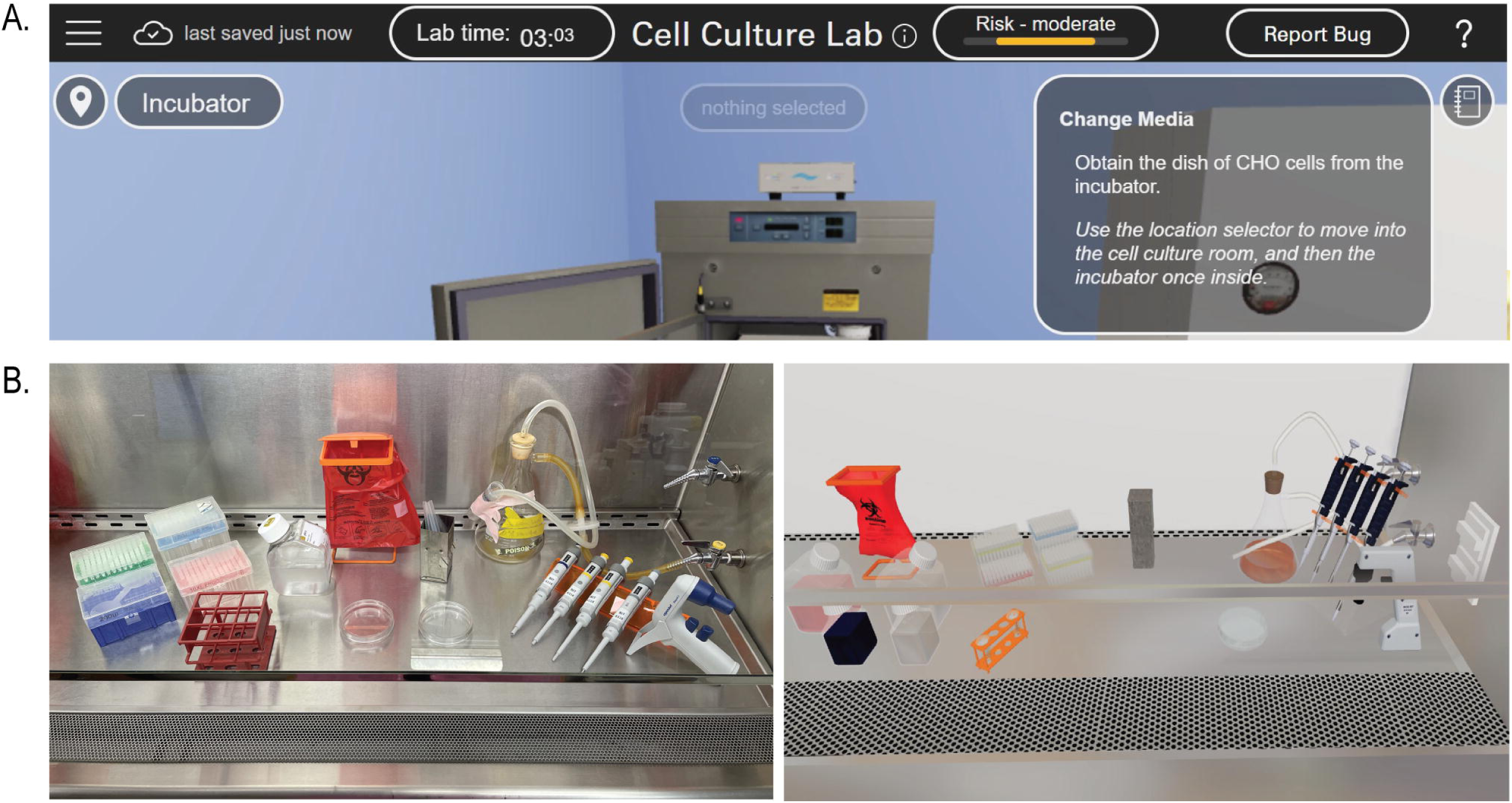
The immersive virtual laboratory cell culture simulation interface provides real-time feedback and is realistic. (A) The browser-based interface includes a risk-meter that provides students real-time feedback on their laboratory technique. The risk level notification changes as the student performs behaviors that would lead to a contamination risk of their cell cultures. (B) The virtual simulation was modeled off real-world laboratory equipment and facilities. Physical biosafety cabinet (*left panel*); virtual biosafety cabinet (*right panel*).

### Students’ perceptions of the interactive virtual cell culture simulation

#### General perceptions

To address our first research question (determine students’ perceptions of the interactive virtual cell culture simulation), we administered a survey both at the end of the laboratory period and at the end of the semester to determine students’ perceptions of the interactive simulation. Participant perceptions of the simulation remained stable during these time periods, with paired sample *t*-tests revealing no significant differences between time points (**S2 Table**). Overall, participant perceptions were positive in regards to the simulation requiring them to think critically, helping them make connections between prior and new knowledge, increasing their understanding of the importance of sterile mammalian cell culture technique, and providing real-world applications (**Fig 2 and S2 Table**). However, participants reported more neutral perceptions of the simulation increasing their engagement (**Fig 2**).

**Fig 2.**
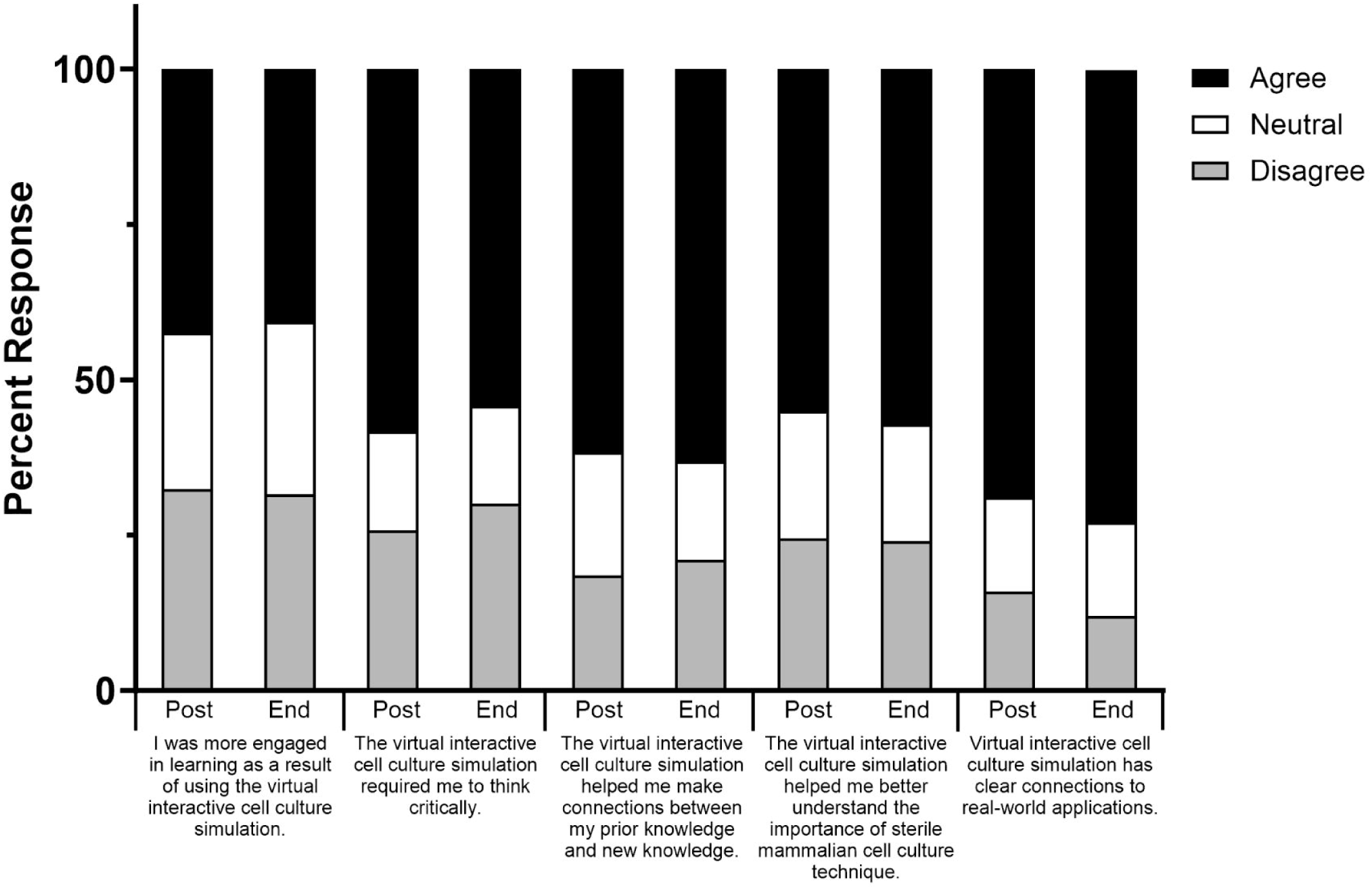
General student perceptions towards the interactive virtual cell culture simulation. Student perceptions were determined both immediately after using the virtual cell culture simulation (post; n=151) and at the end of the course (end; n=134) using an online survey. Values indicate the overall percentage of student responses for each specified question. *Note:* scale was collapsed across categories for ease of interpretation. Disagree= Somewhat and strongly disagree; Neutral = Neither agree nor disagree; Agree= Somewhat and strongly agree.

#### Thematic Results of Open-Ended Feedback

The coded participant responses fell into six recurring themes among responses to the open ended survey question, “please provide 1-2 things that you have found the most helpful about the virtual interactive cell culture simulation.” **Table 3** provides these themes as well as representative quotes. In total, 43.36% of participants made statements regarding how the simulation promoted their learning, 30.97% made statements regarding the realistic nature of the simulation in comparison to the in-person physical lab, 15.93% made statements on how the simulation helped them better prepare for the physical lab, 15.04% made statements on the helpfulness of the real-time feedback in the virtual simulation, 13.27% made statements regarding the simulations accessibility. Even though it did not meet the criteria for a theme, several participants (n = 5) commented that the simulation provided a training environment that reduced their fear of failure and afforded them opportunities to repeat the protocol multiple times (n=5). This is important since scientific inquiry requires risk taking [41, 42] and competence [25, 43]. However, many participants are adverse to risk-taking behaviors as failure can be associated with risk aversion and a negative stigma [44]. For example, the participant quotes below contain descriptions where they perceive the virtual simulation as a low-risk way to practice their biotechnology skills.

**Table 3.**
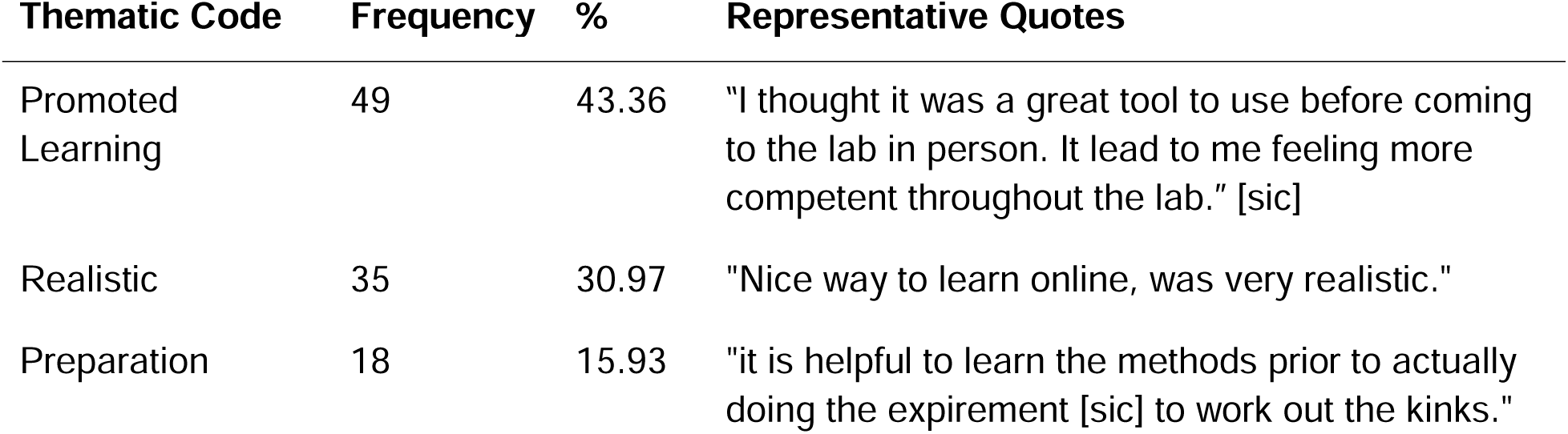
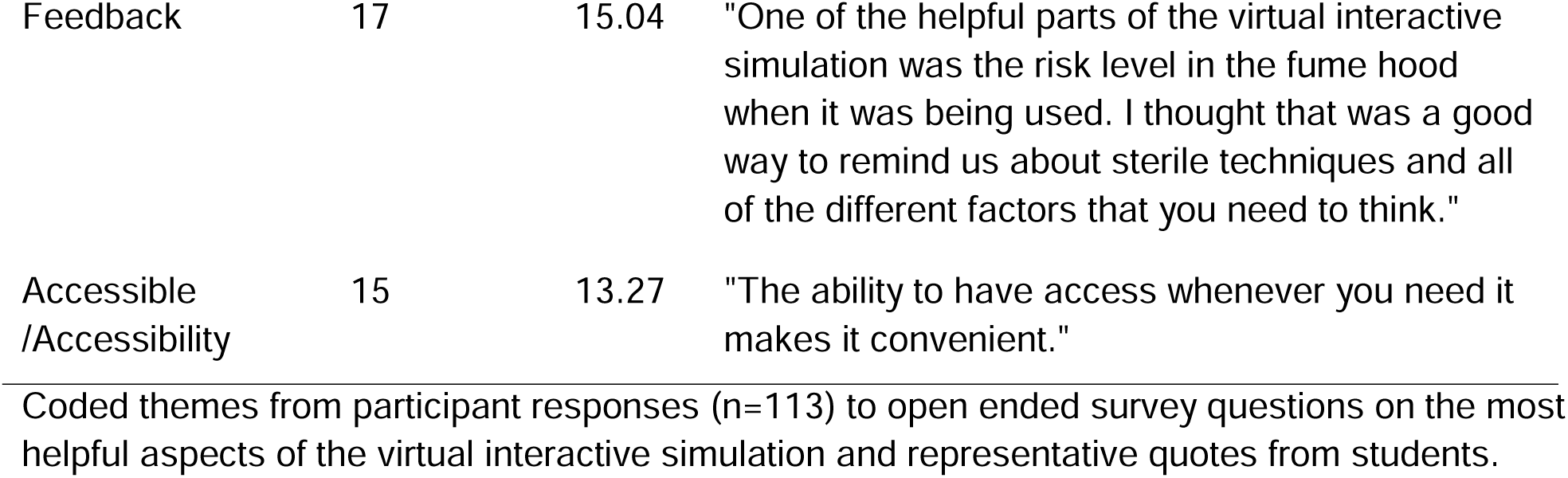
Thematic Codes from Open-ended Student Responses.

Student 1, *“It was nice to see the process in a low-risk way that doesn’t risk ruining precious samples.”*

Student 2, *“I enjoyed the simulation, as it was a great way to practice techniques and using laboratory technology without the cost and any worries about messing up!”*

Student 3*, “*“*It was helpful to have an environment to explore without fear of doing anything incorrectly; occasionally the anxiety of adding improper reagent or performing a step incorrectly can lead to some paralysis during the laboratory session.*”

### Impact of virtual simulation on students’ motivational beliefs

We next wanted to determine if engaging in the virtual interactive cell culture simulation as a teaching tool would impact students’ motivational beliefs regarding biotechnology self-efficacy and science identity. While research has shown that physical laboratory training [45] and virtual lab simulations in microbiology [36] increase students self-efficacy and motivation, less is known about the influence of virtual laboratory training on self-efficacy in biotechnology settings that require complex instrumentation, expensive reagents and advanced protocols. Therefore, to address this second research question, validated survey questions pertaining to self-efficacy [28] and science identity [17] were given to participants at the beginning of the lab period, at the end of the lab period and finally at the end of the semester. Participant responses were then broken down into three groups based on their general perceptions to the simulation (i.e. participants that had high, moderate or low levels of agreement that the simulation was helpful) and analyses conducted to determine the temporal nature of motivation based on simulation perception.

Participant perceptions of the simulation did not impact their beliefs about their ability to complete biotechnology related tasks. No significant timexgroup interaction (F(4, 260) = 1.05, *p* = .38) or main effect of group (F(2, 130) = .98, *p* = .38) were observed. However, the 3x3 (time x group) mixed ANOVA did reveal a main effect of time (F(2, 260) = 10.16, *p* < .001, η^2^ = .07). Bonferroni post hoc tests indicated that participants reported lower perceptions at pre-lab (*M* = 25.04, *SD* = 3.79) compared to post-lab (*M* = 26.18, *SD* = 4.06; *p* < .001) and end of semester (*M* = 25.73, *SD* = 4.24; *p* = .035) (**Table 4**). These data suggest a potential impact of the simulation on self-efficacy. Participants showed significant gains from pre to post lab surveys, with these gains being maintained at the end of the semester (**Table 4**).

**Table 4.**
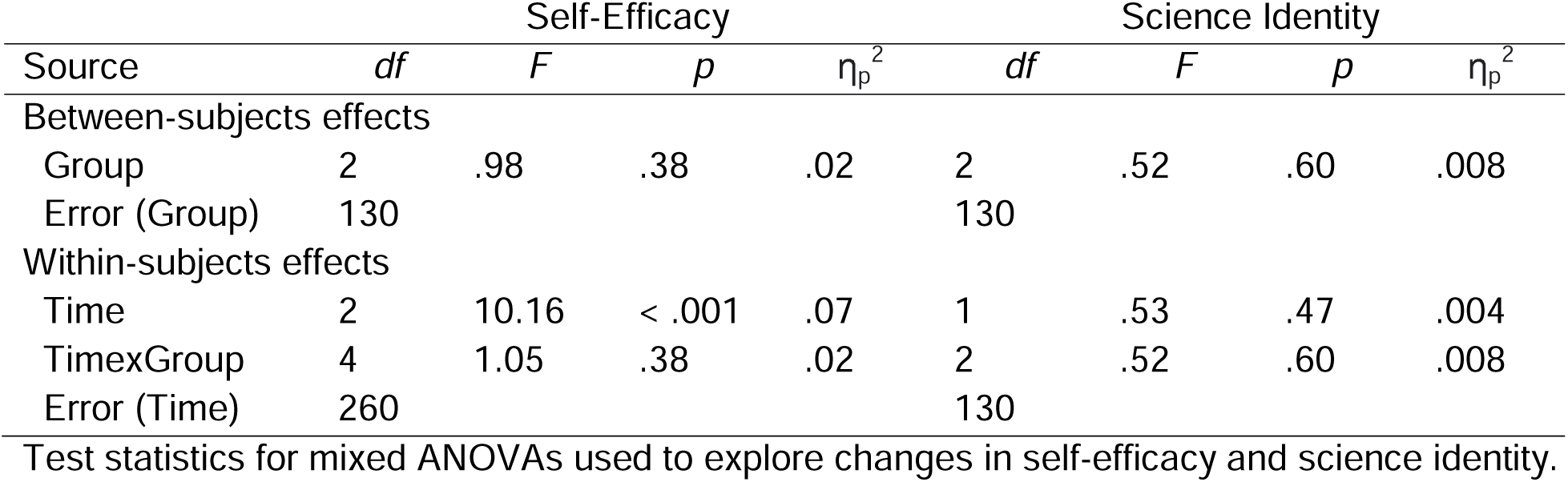
Changes in Student Motivational Beliefs after Using the Virtual Simulation.

Since the effect of virtual laboratory training on science identity has not been explicitly examined in the literature [3], we wanted to determine if engaging in the virtual simulation affected this motivational belief as it is an important metric for student persistence and success in STEM fields [23]. The 2x3 (time x group) mixed ANOVA showed no significant main effect of time (F(1, 130) = .53, *p* = .47), group (F(2, 130) = 2.68, *p* = .07), or timexgroup interaction (F(2, 130) = .52, *p* = .60) (**Table 4**). These data demonstrate that there was no significant effect of the simulation on participants’ beliefs about their science identity. Moreover, engaging with the simulation did not alter participant perceptions/beliefs over time.

## Discussion

Laboratory-based education for students in STEM disciplines is an important component to their knowledge, skills, and professional development. In the last decade, virtual laboratories have been developed to complement or serve as alternatives to traditional physical laboratories. Virtual labs have been positively received by students to promote their learning [6, 7] and have been shown to have comparable learning outcomes compared to traditional pedagogical methods [36]. However, few reports exist on how virtual laboratories may affect students’ belief systems when used in an advanced laboratory-focused interdisciplinary STEM course. In the current study, we investigated student perceptions and motivational beliefs after engaging in an immersive and interactive virtual simulation in an upper-level biotechnology course.

We first wanted to determine students’ perceptions on using the virtual simulation prior to performing a similar laboratory in the physical space. Participants had positive perceptions after engaging in the virtual simulation. In particular, they perceived the simulation as requiring them to think critically, increase their understanding and providing-real world applications and these perceptions remained stable over time even after participating in the physical lab (**Fig 2 and S2 Table**). These perceptions are in alignment with participants’ open-ended feedback. Close to half of participants commented that the virtual simulation promoted their learning, 30% viewed the simulation as realistic and ∼15% commented that the simulation adequately prepared them to learn the techniques, valued the real-time feedback and its accessibility (**Table 3**). Moreover, participants commented that the virtual simulation was a place they could experience failure in a low-risk way, which is important to the learning process as scientific inquiry requires risk taking [41, 42]. On the other hand, participants felt more neutral in their perceptions of the simulation increasing their engagement (**S2 Table**). This may be due to the timing of the simulation as a teaching tool, since it was used immediately prior to students performing the identical physical lab. Overall these data suggest that the virtual simulation is valued by students and can be applied to help them successfully navigate physical laboratory techniques.

Instructors’ observations indicate students that completed the virtual laboratory simulation prior to the physical lab were more prepared, made less mistakes and completed the lab in less time than students that did not engage in the virtual simulation ahead of time (unpublished observations). In addition, the majority of students met the learning outcomes for the lab concepts as assessed through submission of an electronic lab notebook entry (data not shown). However given the nature of the assessment, we were unable to directly determine the effectiveness of the virtual simulation alone in bolstering biotechnology technical skills. Therefore, future work will further develop simulation analytics to evaluate how virtual behaviors and lab outcomes correlate to physical outcomes. Moreover, we are assessing the educational impact of biotechnology skills training with the virtual simulation alone (i.e. distance education courses) compared to students who have access to both virtual and in person training.

In addition to student perceptions, we also evaluated student motivational beliefs. While a number of studies have evaluated the benefit of virtual simulations in increasing the cognitive domain of learning (i.e. specific knowledge, skills) [2] few have addressed the effects on the affective domain especially in biotechnology education [36, 46]. Interestingly, participant perceptions of the simulation did not influence their biotechnology self-efficacy (**Table 4**). In fact, all participants regardless of their overall perception of the simulation demonstrated a significant increase in self-efficacy from the pre to post course survey (**Table 4**). The increase in self-efficacy scores of participants may suggest that the virtual simulation is a valuable tool to help students perform experiments by affording them the ability to iterate and master skills thus increasing their confidence [31, 33]. However, it should be noted that participants engaged in both the virtual simulation and the physical laboratory prior to completing the post course survey. Therefore, the increase in self-efficacy could potentially be attributed to both or one of the training modalities. Importantly, the use of the virtual simulation in this context did not deter but rather bolstered participant beliefs that they can complete biotechnology-related tasks.

A similarly important belief correlated with self-efficacy is science identity. Participants exhibited a high science identity prior to engaging in the virtual simulation and their science identity scores did not significantly change throughout the duration of the course (**Table 4**). The virtual simulation was intentionally designed to include elements that foster a positive science identity akin to their experiences in the physical lab as students apply their technical skills through virtual hands-on experiences [25]. These data suggest that engaging with the simulation and a physical lab does not alter their scientific identity. We did note that the majority of participants who responded to the survey had high science identity beliefs. This is likely due to the nature of the course, which is geared towards upper level STEM undergraduate and graduate students. The course is not required for any major and many students opt to take the course based on the content’s relevance to their career and research aspirations. Therefore the participants in this study may be self-selected as those that intrinsically have a higher science identity having persisted in their science-related majors (**Table 1**) [15, 47, 48].

Collectively, our data suggest that using a virtual simulation as a training tool in an upper level biotechnology course was positively perceived as a benefit to participants by requiring them to think critically, practice their technical skills in a real-world application and support their identity as scientists. Furthermore, participants demonstrated an increase in self-efficacy when the virtual and physical laboratories were paired together. Therefore the virtual simulation can be a useful pedagogical tool that lowers accessibility barriers of a physical lab, offers opportunities for students to iterate techniques and is easy to implement.

## Acknowledgements

The authors would like to thank the students enrolled in the course for their feedback and participation. This work was supported through funding by a NC State DELTA Exploratory Grant and BioMADE Education and Workforce Development Grant [FA8650-21-2-5028] to M.C.S. The authors would also like to thank the NC State Biotechnology Program for their support.

## Supporting information

**S1 Table.** Cronbach’s alpha (α) measure of scale reliability for biotechnology self-efficacy and science identity.

**S2 Table.** Comparisons of perceptions towards the virtual interactive cell culture simulation post lab and end of semester.

## Notes

### Competing Interest Statement

The authors have declared no competing interest.

