## Supporting Tables 1 and 2 for "Student experiences with an interactive 3D immersive biotechnology simulation and its impact on motivational beliefs"

### **Supporting Information**

**S1** **Table.** Cronbach’s alpha (α) measure of scale reliability for biotechnology self-efficacy and science identity.

|  | Survey | | |
| --- | --- | --- | --- |
| Measure | Pre-Lab | Post-Lab | End of semester |
| Biotechnology Self-Efficacy | .84 | .90 | .89 |
| Science Identity | .88 | - | .94 |

**S2** **Table.** Comparisons of perceptions towards the virtual interactive cell culture simulation post lab and end of semester.

| **Item** | **Post-Lab** | **End of Semester** | **t-test** |
| --- | --- | --- | --- |
| I was more engaged in learning as a result of using the virtual interactive cell culture simulation. | 3.07 (1.21) | 3.09 (1.09) | t(131) = -.23, p = .41 |
| The virtual interactive cell culture simulation required me to think critically. | 3.44 (1.25) | 3.33 (1.20) | *t*(131) = 1.16, p = .12 |
| The virtual interactive cell culture simulation helped me make connections between my prior knowledge and new knowledge. | 3.60 (1.11) | 3.52 (1.10) | *t*(131) = .86, p = .20 |
| The virtual interactive cell culture simulation helped me better understand the importance of sterile mammalian cell culture technique | 3.50 (1.22) | 3.45 (1.21) | *t*(131) = .42, p = .34 |
| Virtual interactive cell culture simulation has clear connections to real-world applications | 3.73 (1.08) | 3.80 (1.04) | *t*(131) = -.71, p = .24 |
| *Note:* post-lab and end of semester columns include mean and (standard deviation) | | | |
